# Seizures initiate in zones of relative hyperexcitation in a zebrafish epilepsy model

**DOI:** 10.1101/2021.03.30.437750

**Authors:** James E. Niemeyer, Poornima Gadamsetty, Chanwoo Chun, Sherika Sylvester, Jacob P. Lucas, Hongtao Ma, Theodore H. Schwartz, Emre Aksay

## Abstract

Seizures are thought to arise from an imbalance of excitatory and inhibitory neuronal activity. While most classical studies suggest excessive excitatory neural activity plays a generative role, some recent findings challenge this view and instead argue that excessive activity in inhibitory neurons initiates seizures. We investigated this question of imbalance in a zebrafish seizure model with multi-regional two-photon imaging of excitatory and inhibitory neuronal activity using a nuclear-localized calcium sensor. We found that seizures consistently initiated in circumscribed zones of the midbrain before propagating to other brain regions. Excitatory neurons were both more prevalent and more likely to be recruited than inhibitory neurons in initiation as compared with propagation zones. These findings support a mechanistic picture whereby seizures initiate in a region of hyper-excitation, then propagate more broadly once inhibitory restraint in the surround is overcome.

**Teaser:** We uncover the roles of excitation and inhibition during seizures, thus opening a path to more targeted therapy of epilepsy.

## MAIN TEXT

### Introduction

Excitation and inhibition are countervailing forces of brain activity that, in normal brains, operate in balance. However, in certain pathological brain states, such as epileptic seizures, this excitation:inhibition (E:I) balance goes awry. While traditional doctrine held that seizures resulted from hyperactive excitatory drive (1–4), recent studies in various models have suggested a generative role for inhibitory neuronal subtypes in ictal onset. This revised picture has been supported by observations of increased inhibitory activity in pre-seizure periods (5–8). For example, low-voltage fast seizures in humans are marked by early inhibitory cell firing prior to peak excitatory cell activity (7). Evidence for a direct causal role of inhibitory cells in seizure onset is also reported (9–11). For example, blocking the effects of inhibitory activity with a GABA antagonist (picrotoxin) can paradoxically prevent spontaneous seizures in a genetic mouse model of human autosomal-dominant frontal lobe epilepsy (9). Another study showed that optogenetic stimulation of a specific interneuron subtypes (parvalbumin- or somatostatin-positive cells) can initiate seizures (11). Together, these finding challenge the traditional concept of excitatory neuron hyperactivity as the primary driver of seizure initiation.

However, the role of inhibitory cells in seizure onset is still widely debated. Not only is there data showing that inhibitory cell activation can inhibit seizure onset (12), but activation of inhibitory cells has been clearly shown to shorten seizure duration (13, 14). While differences in seizure models, experimental preparations, and recording techniques could explain these disparate results, the explanations may be more subtle. While one recent study found that inhibitory cells can exert either anti- or pro-seizure effects depending on the timing of activation (15), another showed that the effects of excitatory and inhibitory cells may be spatially dependent (16). These findings suggest that the role of E:I balance in seizures may not be as simple as “more E” or “earlier I”, but rather that the effects of E:I balance on ictal onset and evolution may be spatiotemporally contingent.

Understanding how E:I balance plays into ictogenesis has several experimental requirements. First, a large region of brain, inclusive of various interconnected structures, must be analyzed during seizure periods. Ideally, this areal sampling would include both the site of seizure initiation as well as sites of later seizure propagation. Second, the neural activity analyzed should simultaneously include both E and I information that can be separated with single-cell resolution. Finally, the neural activity should be recorded in awake animals whose activity can best recapitulate seizures lacking the confounding effects of anesthesia.

These criteria can be relatively easily met in the zebrafish preparation (17). These vertebrates exhibit significant homology with humans at the genetic (18), neurochemical, and brain structural (19) levels. Their transparency, rapid development, and small size in the first few weeks after fertilization allows whole brain optical access at sub-cellular resolution throughout the brain. Their genetic tractability permits tissue and cell-type specific experimentation. Further, their ease-of-handling facilitates both high-throughput drug screening (20) and high-resolution imaging and electrophysiological studies in the unanesthetized animal. Prior work on seizures using this animal model has already shown similar electrographic profiles (21), glial involvement (22), and responses to seizure treatments (23) as found in human epilepsy.

Here we used the zebrafish model to investigate spatiotemporal variations in E:I balance during seizure initiation and propagation. We monitored seizure dynamics using both electrophysiological recording and two-photon calcium imaging that enabled tracking of neuronal dynamics across the brain at the single cell level. We found specific regions of the brain consistently associated with seizure initiation, while others with seizure propagation. Using transgenic specification to separate excitatory and putative inhibitory populations, we found that the ratio of E:I activation at ictal onset was greatest in initiation zones. This finding demonstrates the importance of early excitation at triggering ictal events in the zebrafish model and challenges recent reports arguing that early inhibitory neuron activity is always the key driver in initiating seizures.

### Results

We present our investigation of the spatiotemporal dynamics of E:I balance during seizures as follows. First, we introduce the experimental preparation and describe electrophysiological characterization of inter-ictal and ictal events in this epilepsy model (**Fig. 1**). Next, we present a pixel-based timing analysis (**Fig. 2**) that is used to create maps of the spatiotemporal dynamics in the brain during seizures (**Fig. 3**). We then examine timing at a cellular level in excitatory and inhibitory populations in regions designated as ‘initiation’ and ‘propagation’ zones (**Fig. 4**). Finally, we confirm that findings from our single-cell analysis hold at the network level (**Fig. 5**) and when examined with spatial specificity using correlation analysis (**Fig. 6**).

**Figure 1:**
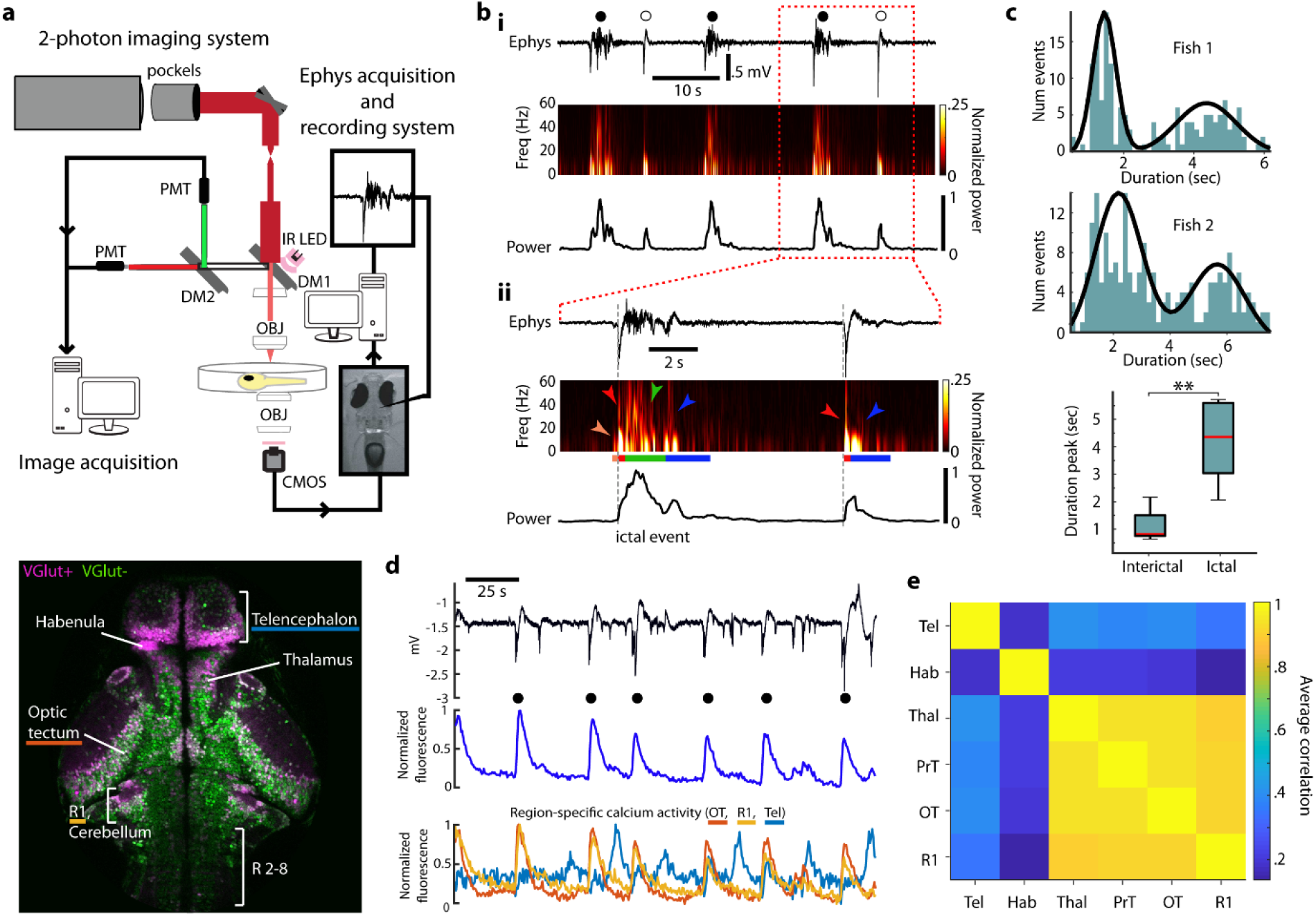
Ictal and interictal activity in the larval zebrafish. (**A**) Top: experimental setup for 2-photon imaging and electrophysiology in larval zebrafish during seizures. Bottom: single imaging plane revealing distribution of VGlut+ (magenta) and VGlut- (green) cells in a variety of brain regions (R: rhombomere). (**B**) Electrophysiological signatures of ictal- and interictal-like events in the zebrafish. (i) Top: Local Field Potential (LFP, top); Middle: spectrogram; Bottom: total power during a period of ictal (closed circle) and interictal (open circle) events; (ii) Similar organization as above but at finer time resolution, with indication of distinct sections (orange, red, green, blue) characterizing the time-course of the two types of events. (**C**) Top, Middle: histogram of event duration times for two fish (green) and best fit (black) with a gaussian mixture model; Bottom: for all fish, duration time for ictal and interictal events. (**D**) Top: LFP, Middle: global change in GCaMP6f fluorescence; Bottom: regional change in fluorescence during a series of ictal (closed circle) and interictal events (OT, optic tectum; R1, rhombomere 1; Tel, telencephalon). (**E**) Matrix of correlations in the activity among various brain regions (Tel: telencephalon, Hab: habenula, Thal: thalamus, PrT: pretectum, OT: optic tectum, R1 rhombomere 1).

**Figure 2.**
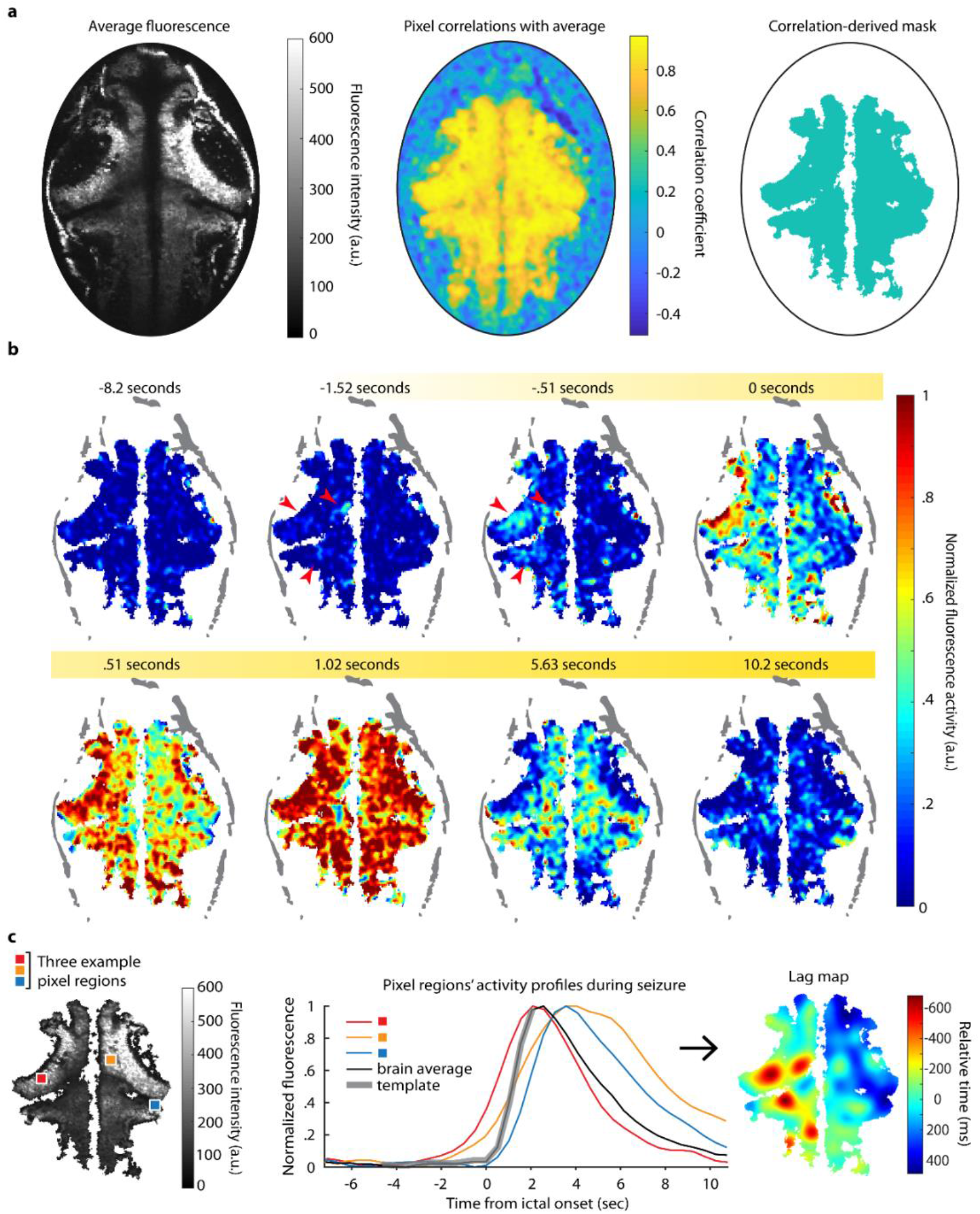
Ictal event mapping. (**A**) Left: average fluorescence over a series of ictal events in one brain; Middle: pixel-wise correlation with brain average; Right: the correlation mask. (**B**) A single seizure’s fluorescence activity is shown over time; red arrows highlight early activity. (**C**) Left: sample pixels and their activity profiles (*middle*) over a single seizure are shown; Right: the corresponding smoothed lag map for this seizure shows the relative activation times for each pixel with respect to the brain average.

**Figure 3.**
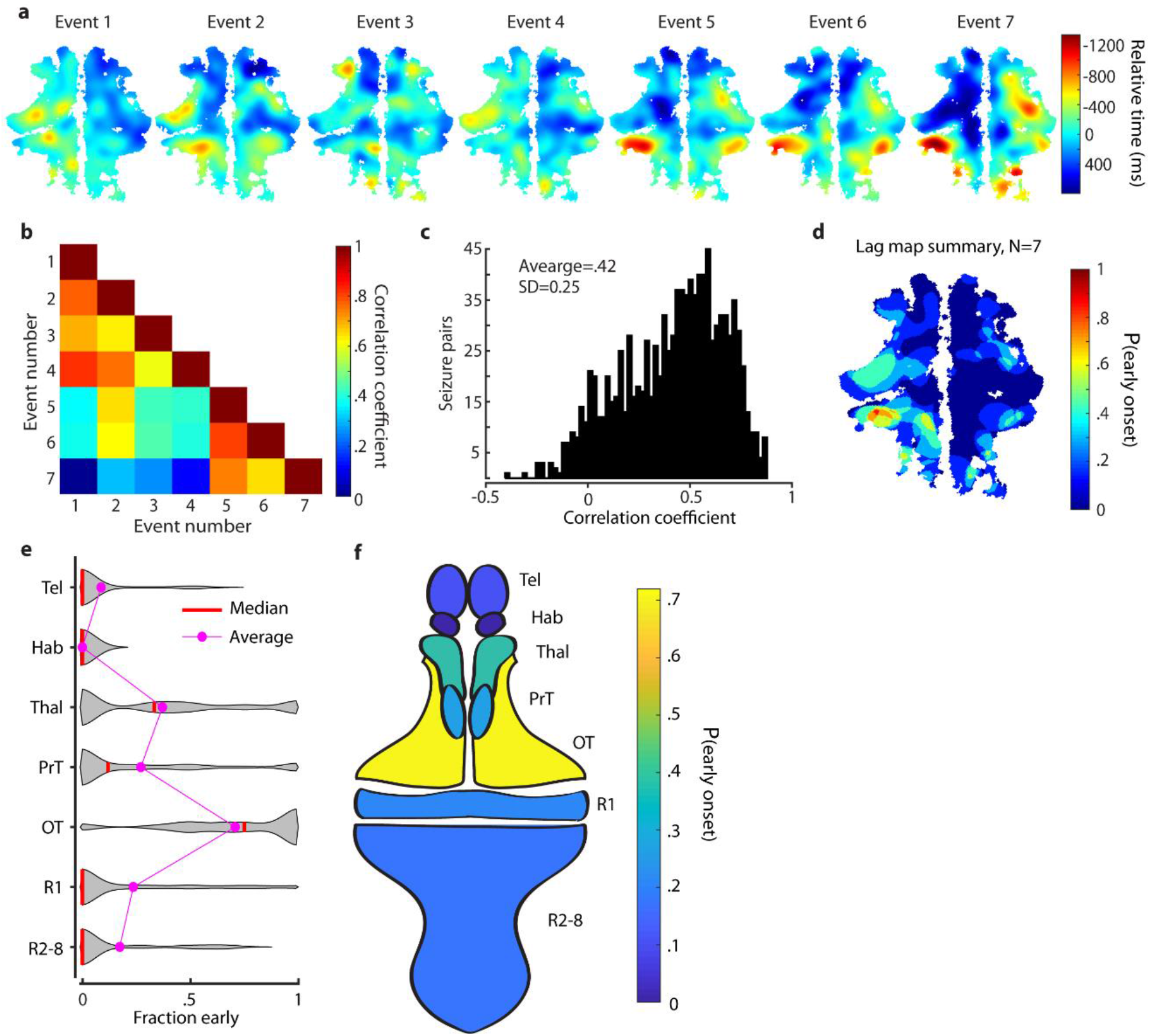
Ictal events evolve similarly within fish, but initiate in different regions across fish. (**A**) Lag maps for seven seizures in one animal. (**B**) Correlations between the events in (**A**). (**C**) correlations of all pairs of ictal events recorded. (**D**) A summary map of the ictal events in (**A**). Violin plots (**E**) and a schematic fish brain map (**F**) of early-onset regions show where ictal events initiated most commonly across different fish.

**Figure 4.**
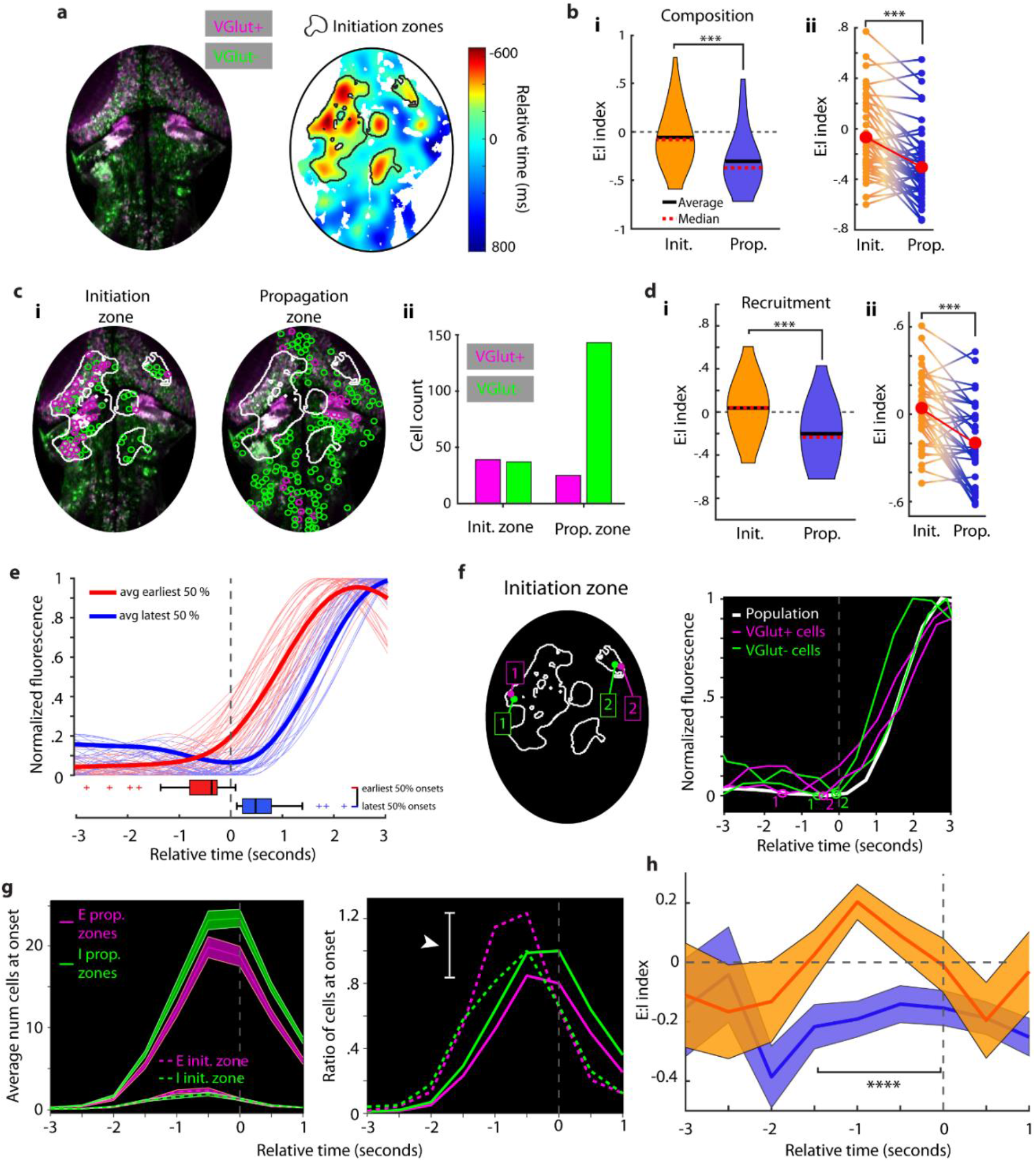
Ictal events preferentially initiate in excitatory cell-rich regions of the zebrafish brain. (**A**) A sample fish image with E and I cells differentially labelled and its associated lag map and initiation zone for a single ictal event analyzed in this figure. (**B**) E:I index in initiation and propagation zones (i) and within-seizure E:I comparisons (ii). (**C**) Single cell locations in a seizure, split by initiation vs propagation zone location (i) and cell counts from initiation and propagation zones (ii). (**D**) E:I index of cells recruited to seizures in initiation and propagation zones (i) and within-seizure E:I index comparisons. (**E**) Individual cell activity traces (smoothed and interpolated for clarity) are shown for one ictal event. Color groups denote onset times from the earliest 50% (red) and latest 50% (blue) of onsets. Boxplots show distributions of onset times. (**F**) Left: sample excitatory and inhibitory cells from initiation zones; Right: their associated calcium traces during this event. Detected onset times are plotted on traces. (**G**) Left: average number of cells at onset across all ictal events grouped by initiation or propagation zone membership; Right: the ratio of cells at onset (E:I) over time, normalized to by the number of inhibitory cells at onset. (**H**) E:I ratio index calculated in initiation (orange) and propagation (blue) zones. All error bars are mean +/- SEM.

**Figure 5.**
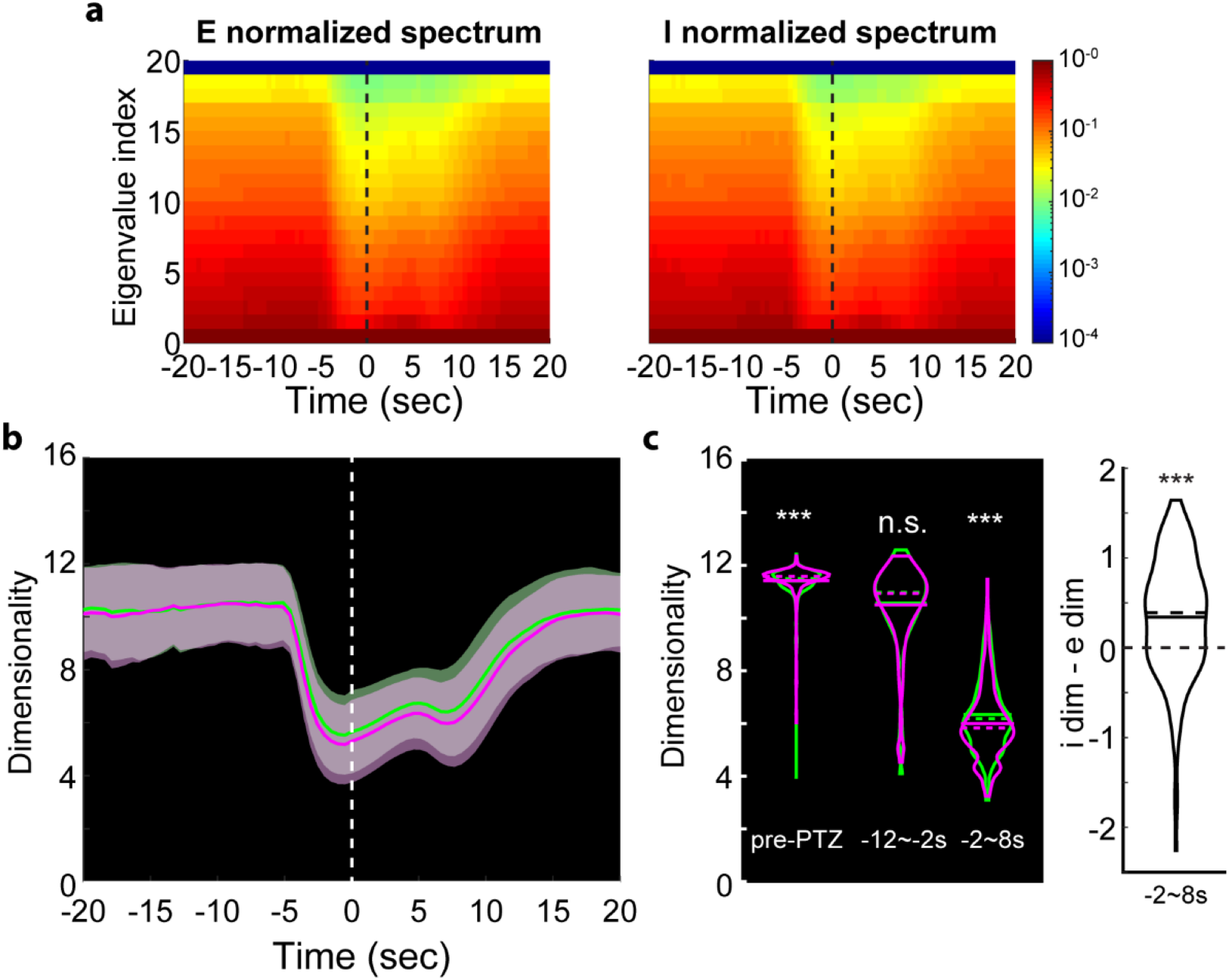
Dimensionality of the excitatory and inhibitory cell population activities. (**A**) Left: Eigenspectrum of the excitatory population activity over 10 sec.-wide rolling window, normalized to the greatest eigenvalue per time point. Vertical line at time 0 is the ictal onset point. Right: same analysis done for inhibitory cell population. (**B**) Average trends of the dimensionality. Vertical line at time 0 is the average ictal onset point. Dimensionality over 10 sec moving window (magenta: e-e; green: i-i). (**C**) Left: Dimensionalities of i-i pairs and e-e pairs shown separately for pre-PTZ, −12~-2 second and −2~8 second periods relative to the average ictal onset point. Right: differences in I-E dimensionalities at −2~8 seconds.

**Figure 6.**
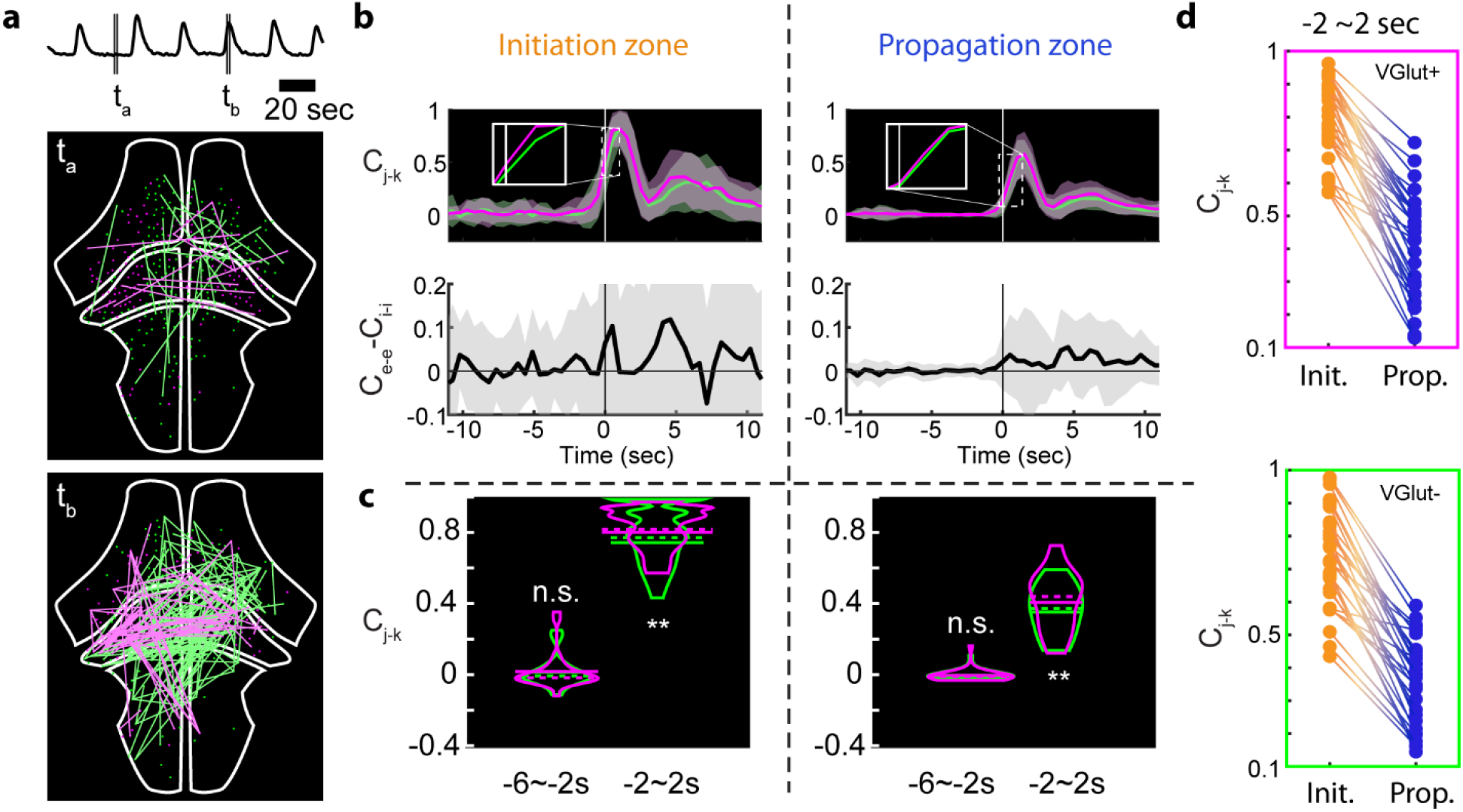
Excitatory and inhibitory cell network correlations. (**A**) Visualization of activity correlations at the cross-section of the whole brain. Pairs with correlation values higher than 0.999 are shown. Top: average normalized fluorescence trace. Middle: correlations during interictal period. Magenta: highly correlated e-e; Green: i-i. Bottom: correlations during ictal period. (**B**) Average trends of interictal to ictal transition for the initiation and propagation zones. Top: correlation metric calculated over 1.5 sec.-wide rolling window (magenta: e-e; green: i-i). Bottom: difference of e-e and i-i correlations; (**C**) Correlation metric values of i-i pairs and e-e pairs shown separately for −6~-2 second and −2~2 second periods relative to the average ictal onset point. (**D**) Within-seizure correlations of e-e cell pairs (top) and i-i cell pairs (bottom) between initiation and propagation zones.

#### Experimental paradigm for examining excitation and inhibition

Experiments were performed using larval zebrafish with all neurons expressing GCaMP6f and VGlut2 cells expressing DsRed (**Fig. 1A**). The first fluorophore, H2B-tethered GCaMP6f, is a fast nuclear-localized calcium sensor that is localized to the nucleus to eliminate contamination from neuropil signal and enable better separation of signals between cells (24). The second fluorophore, DsRed, was used to help separate excitatory from inhibitory neurons. VGlut2 is a vesicular glutamate transporter gene found in the vast majority of glutamatergic cells in the larval zebrafish (25). The largest proportion of VGlut2-cells are inhibitory GABAergic neurons, with non-trivial contribution from excitatory VGlut1 cells (primarily cerebellar granule neurons) and inhibitory glycinergic neurons. Due to the purely excitatory nature of the VGlut2 population and the primarily inhibitory nature of the VGlut2-populations, below we refer to the former as ‘excitatory (E)’ neurons and the latter as ‘inhibitory (I)’ neurons.

#### Seizure characterization by electrophysiology and calcium imaging

Our first goal was to characterize seizures measured electrographically. Seizures were induced using bath applied PTZ in larval zebrafish that were held in agar. While the definition of a “seizure” can be flexible and the exact features of seizures likely vary between species and models, our electrophysiology data below demonstrates separable ictal-like and inter-ictal-like events with electrographic signatures similar to those observed in other animal models and humans. For simplicity, here we use the word “seizure” to refer to the longer ictal events as others have done for PTZ-induced ictal activity in zebrafish (22, 26, 27). In initial two-electrode experiments, we placed electrodes in different brain regions (telencephalon, optic tectum, or hindbrain) and found that ictal events developed concurrently after ~15 minutes of PTZ application. Electrodes placed in the optic tectum consistently captured the largest set of ictal events (unpublished observations)—hence, in subsequent experiments, we used only one electrode placed in the neuropil of the optic tectum. **Figure 1B** shows an example electrophysiology recording, demonstrating ictal-like events lasting for several seconds (‘ictal’, filled circle symbol), followed by quiescent periods and occasional brief interictal-like events (‘interictal’, open symbol), as well as their corresponding power spectrograms (**Fig. 1B)**. Ictal events (left) were marked by an initial low frequency power increase (orange), a brief cross-frequency spike with most power <20 Hz (red), a sustained phase with a more uniform cross-frequency distribution (often up to 100 Hz) that lasted for several seconds (green), and a recovery phase at lower frequency with gradual relaxation (blue). In contrast, interictal events (right) only exhibited the short cross frequency event (red) before moving to a briefer recovery phase than seen with ictal events (blue). Both forms of event then returned to baseline. While more complicated classification schemes could be used, separation of ictal from interictal events based on duration provided a robust method as ictal events typically lasted 4 times longer than interictal ones (**Fig. 1C**; Wilcoxon signed-rank test, N=9 fish, 1685 events, p=0.003).

We next employed simultaneous cellular-resolution 2-photon imaging of larval zebrafish along with the electrophysiological recordings. Ictal events were clearly observable in population activity and within specific brain regions (**Fig. 1D**; **Fig. S1**). Interictal events, on the other hand, did not lead to global changes in activity. Examining global activity spatially, we found that ictal events did not typically initiate in or recruit the telencephalon (**Fig. 1E**). While activity in the optic tectum (OT) and rhombomere 1 (R1) correlate strongly with the whole-brain average calcium response, the telencephalon (Tel) does not. A correlation matrix built with average calcium activity in different brain regions from multiple series of seizures from different fish (N=5 fish, 22 seizures) indicates that activity in the telencephalon and habenula is largely uncorrelated from the rest of the brain during ictal events.

#### Lag maps identify regions of seizure initiation and illustrate seizure propagation

To examine the involvement and timing of different brain areas in PTZ seizures we next developed a method to extract seizure propagation maps within individual fish. We first used correlation to global average ictal activity to create a mask that excluded pixels where there were no changes in fluorescence related to the seizure (**Fig. 2A**). The activity in the retained pixels was not monolithic, with some regions beginning to show elevations in fluorescence well before other regions (**Fig. 2B,** red arrows). To examine these temporal variations more carefully, we correlated pixel-level activity to that of a template of ictal activity initiation defined from a segment of the global fluorescence event (**Fig. 2C,** *middle*), incrementally shifting the template in time to identify when activity in particular pixel began to rise (see Methods). The template shift times for each pixel defined a propagation map for each ictal event (**Fig. 2C**, *right*). This template matching procedure allowed identification of the earliest portions of the ictal activity in each pixel and was less susceptible to errors than procedures that relied upon threshold crossings or features such as peak fluorescence.

We applied this algorithm to create a lag map for each individual ictal event (N=10 fish, N=177 ictal events). **Figure 3A** shows the lag maps of 7 ictal events in one larval zebrafish (the same animal as **Fig. 2**). This revealed a similarity between neighboring ictal events (e.g., Events 1-4 vs 5-7). In this animal the left cerebellum and the optic tectum showed consistent early involvement in every ictal event. To quantify how similar ictal propagation appeared between ictal events, we measured the correlation coefficient of the 2-D spatiotemporal lag maps of all pairs of ictal events for this animal (**Fig. 3B**). This similarity was quantified for the entire data set by pooling within-animal correlations, revealing a positively skewed distribution (**Fig. 3C,** average Pearson’s *r*=0.42 +/- 0.25, N=177 seizures).

We next developed a scheme to summarize these spatiotemporal patterns and identify regions consistently involved in seizure initiation. Lag maps for individual ictal events were binarized by assigning a value of 1 to all pixels where neuronal activity initiated at least 500 ms in advance of the population event, and a value of 0 to all other pixels. Averaging these mask arrays across all events within a single fish provided a 2-dimensional summary showing where ictal events initiated most frequently within an individual animal (**Fig. 3D**). For the animal whose seizures are displayed in **Figure 3A**, we see that the left cerebellum and portions of the left tectum exhibit the highest probability of involvement in ictal initiation. Across all animals (N=10), the most common initiation zones were the optic tectum, thalamus, pre-tectum, and cerebellum/R1 (**Fig. 3E-F**). Propagation zones were defined as the remainder of the brain to which the seizure spread outside the initiation zone.

#### Excitation and inhibition examined in seizure initiation and propagation zones

Having identified the ictal initiation and propagation zones, we next focused on the issue of E:I balance in these different regions during seizures. We first asked if there were differences between the initiation and propagation zones in the relative numbers of excitatory and inhibitory cells. Taking advantage of the ease with which single neurons can be distinguished with nuclear-localized GCaMP6f, we labelled individual soma using standard computer vision algorithms and separated them into excitatory and inhibitory cells based on the intensity of the DsRed fluorophore (total N=19,434 cells across 9 fish; example event and seizure-recruited cells shown in **Fig. 4A**). An event-specific comparison of the differences between the initiation and propagation regions in composition was performed by calculating for each ictal event a value for an E:I composition index; the index was defined such that a region with only excitatory cells had a composition index value of 1, all inhibitory cells an index of −1, and an equal number of excitatory and inhibitory cells an index of 0. E:I composition in initiation regions was −0.07 +/0.29 (SEM), while E:I composition in propagation regions −0.30 +/- 0.29 (**Fig 4B**; at cutoff requiring at least 10 E and 10 I cells per initiation site, N=49 seizures, Wilcoxon signed-rank test p<0.001). This difference corresponds to a roughly 61% increase in the E:I ratio in the initiation zone relative to the propagation zone calculated over all animals and all seizures. This finding was robust to variation in the minimal number of cells of each type allowed in each region (**Fig. S3**).

We next investigated if there were differences between the initiation and propagation zones in the relative numbers of excitatory and inhibitory cells that were *recruited* into the ictal event. Cells were deemed to be recruited into the seizure if they exhibited of correlation of 0.8 or better with the global ictal event (**Fig. S3**). The locations of such cells during one seizure event are shown in **Figure 4C**, revealing a much larger E:I ratio of recruited cells within the initiation zone than in the propagation zone. To quantify this observation across, we calculated an E:I recruitment index, defined as above but only for recruited cells. E:I recruitment in initiation regions was 0.04 +/- 0.25 (SEM), while E:I recruitment in propagation regions was −0.20 +/0.28, a significant difference (p<.0001, N=167, Wilcoxon signed-rank test; **Fig. 4D** at cutoff requiring 10 E and 10 I cells per initiation site). This corresponds to a roughly 64% increase in E:I ratio in the initiation zone relative to the propagation zone across seizures, and was robust across varying cutoffs of cell counts (**Fig. S3**). Thus, on average there is both a greater fraction of excitatory cells in initiation zones relative to propagation zones, and these excitatory cells are also recruited at a higher fraction within the initiation zone.

We next examined how E:I recruitment during the ictal event varied over time in the initiation and propagation zones. We observed asynchronous single cell onsets during ictal events: **Figure 4E** shows smoothed and interpolated single cell fluorescence traces for one ictal event, colored by initiation time, demonstrating the temporal spread of single cell activity that these seizures typically displayed. As expected, across our data set we observed significantly earlier onsets in the initiation zones compared to propagation zones (**Fig. 4G**, **Fig. S3C**). Interestingly, in the propagation region, the number of newly recruited excitatory cells was always less than the number of newly recruited inhibitory cells (**Fig. 4G,** *left*, solid lines, N=167 seizures). In contrast, in the initiation zone, the number of newly recruited excitatory cells was comparable to the number of newly recruited inhibitory cells at the beginning and end of the ictal event, but was notably higher at a time approximately 1 second ahead of the ictal onset. To see this difference more clearly, we normalized these recruitment curves to the peak of the number of newly recruited inhibitory cells (**Fig. 4G,** *right*), revealing 1) that the number of newly recruited cells peaked earlier in the initiation zone, and 2) a difference of over 50% at peak in the ratios of newly activated excitatory vs inhibitory cells between the initiation and propagation sites (white arrow, **Fig. 4G**). Quantifying these observations using the E:I recruitment index introduced above, we see that significant differences in the pattern of recruitment between the initiation and propagation regions are present in the period −1.5 to 0 seconds before ictal onset (N=36 seizures with at least 10 E and 10 I cells in initiation zone, Wilcoxon signed-rank test p<0.0001, **Fig. 4H**).

#### Network-level analyses reaffirm a privileged role for excitation at seizure initiation sites

We next examined if this privileged position for excitatory neurons during the initiation of seizures was also apparent in higher order relationships between excitatory and inhibitory cells. We began by using Principal Components Analysis to look at the overall degree of coordination in excitatory versus inhibitory populations during the seizure initiation process. For both groups, the number of dimensions needed to explain population dynamics dropped dramatically at the time of ictal onset (**Fig. 5A-B**). This collapse to lower dimensionality, an indication of greater coordination within a population, was most pronounced for the excitatory population (**Fig. 5C**; pre-PTZ N=2781, post-PTZ N=167, right: I-E dimensionality differences, Wilcoxon signed-rank test, *** p<0.001, **p<0.01, n.s. not significant).

To examine this increased coordination of E cells vs I cells with greater temporal specificity and regional context, we next analyzed correlation structure within these cell types by calculating a correlation metric (*C*) in a rolling time window (**Fig. 6A**). We focused on correlations between excitatory-excitatory (E-E) and inhibitory-inhibitory (I-I) cell pairs. Average correlation was generally low for both pairings during baseline conditions and interictal periods (**Fig. 6B**, *top*). Correlation increased dramatically for both pairings around the time of ictal initiation, with correlation change in the initiation zone on average larger and more advanced in time than that in the propagation zone. We found that E-E correlation was slightly elevated relative to I-I correlation during ictal onset in both regions (**Fig. 6B**, insets); this elevation was clearer when looking at the correlation difference, which was largest around the time of ictal onset in the initiation zone (**Fig. 6B-C,** Wilcoxon signed-rank test, N=36 seizures with 10 E and 10 I cells in initiation zones, *** p<0.001, ** p<0.01, n.s. not significant; **Fig. 6D**, within-seizure correlation differences in initiation and propagation sites in events with 10 E and 10 I cells). Thus, these correlation patterns further suggest that the coordinated activation of excitatory neurons in the initiation zone plays an important role in triggering ictal events.

### Discussion

In this study, we addressed the question of E:I balance in seizure initiation and propagation using the zebrafish model. The main technical limitation afflicting prior studies of this topic has generally been the need for broad, cell-specific sampling of the widespread neuronal substrate inclusive of the ictal onset zone and all areas of potential propagation. We overcame these challenges using a global imaging strategy in the larval zebrafish, an optically transparent vertebrate preparation allowing for simultaneous imaging of multiple cell types. We demonstrated a privileged role for excitation over inhibition in the seizure initiation zone, while regions that were invaded later were biased towards stronger inhibitory cell recruitment.

#### Seizure dynamics in the larval zebrafish

One specific relevance of our findings will be to aid efforts to identify seizure medications with high-throughput screening. Zebrafish are increasingly used to test potential drug treatments for diseases such as epilepsy (23, 28). High-throughput drug screening is a primary method of therapeutic discovery (29), and the zebrafish offers significant advantages in this regard due to the ease with which drugs can be delivered uniformly to the brain while behavioral changes are monitored *en masse*. However, while such high-throughput studies can be quite useful in identifying promising classes of drugs, optimization of both the therapeutic and its dosing will benefit greatly from a mechanistic understanding of the pathophysiology being addressed. The larval zebrafish affords several advantages for understanding the cellular and circuit mechanisms that have been used to study the normal brain, including high-resolution optical access globally, a suite of genetic tools for monitoring and manipulating specific cell types, and the capacity to study all constitutive elements of an intact circuit. Here we have harnessed some of this power to begin understanding the pathophysiology of seizures. The preponderance of excitatory activity in the initiation zone points towards exploiting this model to screen, with greater granularity, classes of drugs that can preferentially regulate the activity of glutamatergic neurons in midbrain structures.

We triggered seizure events in the zebrafish model using PTZ, a chemoconvulsant commonly used in therapeutic screening. PTZ is the only pharmacological seizure-inducing drug employed at the initial identification stage by the NINDS Epilepsy Therapy Screening Program (30). PTZ is known to induce seizures in humans (31, 32) and lower seizure threshold in epileptic patients (33). PTZ-induced seizures in the zebrafish occurred spontaneously, repeatedly, and throughout the brain, consistent with earlier reports (22, 26, 34, 35). Furthermore, the ictal LFP had spectral features (36) similar to those observed in rodent models (37) and human patients (38): at initiation, ictal events displayed an abrupt broad-frequency power increase, followed by a longer period where power was restricted to a slightly lower frequency range, concluding with a return to baseline power at the time of termination. This commonality supports the idea that the PTZ model of ictogenesis in the zebrafish is of general relevance for the study of seizures in other species.

#### Spatio-temporal consistency of seizure initiation and propagation

Regions of ictal onset and propagation were generally conserved within fish, but somewhat varied between animals. Rodent slice experiments have shown that the seizure foci can change slowly over time: one focus can dominate for several minutes before being replaced by another region (39). However, such studies have also found that certain brain regions (e.g., hippocampal CA2-3) and cell layers (neocortical layer 5 pyramidal cells) are predisposed to act as pacemakers in seizure-generating activity (40, 41). Our findings extend these results to an awake, whole-brain model. We found a significant level of consistency in initiation and propagation patterns within fish, although correlations weakened over tens of minutes. While there was some variation between animals, we found that midbrain structures had a quite high propensity to serve as initiation zones. We also confirmed the surprising finding that the telencephalon (homologous with human cortex) was one of the last regions to be recruited in the seizing zebrafish brain (22). Thus, our results provide further evidence that certain subcortical sites may serve a prominent role in generalized seizure initiation (42–45) though these results do not negate a role for cortical contributions.

#### E:I balance differs in initiation and propagation zones

Synchronous excitation has long been thought to drive epileptic seizures (3), while recent cell-specific studies have suggested a role for inhibitory neurons in both seizure initiation (6, 7) and termination (12, 14). Our finding adds to the modern E:I discussion by reaffirming the classic distinction between ictal initiation and propagation. Specifically, we observed a higher ratio of recruited excitatory:inhibitory cells near the seizure initiation site compared to locations that were recruited later by the same seizure. A recent study in anesthetized mice reported a similar finding: the E:I activity ratio, calculated on averaged parvalbumin-positive (PV) and non-PV interneuron activity, was higher in the initiation zone compared to propagation zones during the pre-seizure period (16). However, the experiment involved focal injection of 4-AP, where the assumption that the injection and onset sites are superimposed may not always be correct. Our experiments circumvent this ambiguity by allowing seizures to develop spontaneously in anatomic regions with lower threshold in a model that permits whole-brain imaging so that we could identify distinct initiation zones on an event-by-event basis.

Some studies in rodents (6, 13) and humans (7) have reported earlier activity from inhibitory cell populations before recruitment to seizure events. We did not observe significantly different onset timing between E and I cell types. We speculate that this could be due to differences in recording location (initiation vs propagation zones), or possibly due to variations in technique. For example, Khoshkoo et al. used an optogenetic kindling model (13), and the firing rate changes between inhibitory and excitatory cells observed by Miri et al. were on the order of just ~2 Hz (6), which our calcium imaging may not have been sufficiently sensitive to detect. Our findings therefore do not rule out the possibility of earlier inhibitory cell activation, but do support a model of ictogenesis that depends on higher-weighted excitatory cell recruitment at initiation sites.

Numerous studies have shown that surround inhibition, present outside of an ictal focus, can play a role in restraining seizures (16, 46–50). This restraining penumbra is a region characterized by massive synaptic barrages, but little neuronal firing, until a temporal point at which the seizure annexes this region (16, 47, 49–51). In our study using a nuclear-localized GCaMP to focus purely on somatic activity, we did not observe significant differences in E and I cell timing, but did observe that seizure initiation zones were heavily weighted towards excitation (higher E:I) while later propagation regions were weighted towards inhibition (lower E:I). In this drug-bathed model, initiation zones are selected based on inherent increased E:I whereas propagation zones transiently resist rapid spread, presumably through a mechanism based on higher relative local inhibition. Future work in this model may shed further light on this inhibitory restraint by combining imaging and intracellular recording (50), allowing simultaneous measurement of synaptic barrages, spiking, and whole-brain activity during seizure dynamics.

#### Limitations

The larval zebrafish model is increasingly being employed to test pharmacological drugs and as a model organism of various neurological disorders. However, the direct correspondence of seizure mechanisms in the zebrafish and humans is difficult to determine. Thus, while our finding that seizure initiation is associated with hyperexcitation does align with many previous studies in mammals and humans, our data cannot state certainly that human seizures follow this principle. Similarly, our use of PTZ to induce seizures may also create difficulty in comparing results with human epilepsy patients whose seizures initiate spontaneously without any pharmacological induction. We note, however, that our findings using PTZ will be directly applicable to drug screening experiments, which typically use PTZ to induce seizures. Future experiments using human brain organoids will be enlightening for both of these limitations.

Finally, our data is also limited in cell-type resolution: we separated excitatory and inhibitory cell classes based on the presence of VGlut2a (also called VGlut2.1), a vesicular glutamate transporter. VGlut2b is co-expressed with VGlut2a, but it should be noted that VGlut1 cells also exist in the zebrafish brain, though to a vastly smaller degree (25). Because we infer inhibitory cell identity by lack of VGlut2 expression, some cells marked as inhibitory in our data could be glutamatergic. However, our primary finding is that E:I balance is higher at seizure initiation sites, and if a subset of inhibitory cells were mislabeled across the brain this finding is not likely to be affected.

The goal of epilepsy research is to understand how normal brain activity transitions to an ictal state and how the event spreads across adjacent normal brain regions. In our model we found that midbrain and diencephalic regions were the most common sites of seizure initiation. Further, during individual seizures we observed significantly higher E:I ratios in initiation compared with propagation zones, supporting a model of seizures in which excitatory activity overruns inhibitory restraint to initiate the seizure while distal regions remain more protected by an inhibitory veto (47, 48, 51). The differential E:I balance we report across brain regions provides a framework for designing and interpreting drug screening studies. Further, our finding of hyperexcitation in initiation zones could be leveraged to perturb neural activity in a cell-specific manner to attenuate seizure initiation or even arrest ongoing seizures.

### Materials and Methods

#### Experimental Design

We performed cellular resolution imaging and electrophysiology experiments in awake larval zebrafish undergoing seizures induced by bath application of pentylenetetrazol (PTZ). Animals expressed fluorescent proteins to allow dissociation of excitatory and inhibitory neuron subtypes. We examined regions throughout the brain simultaneously, measuring where seizures in these animals initiated and how excitatory and inhibitory neurons were recruited to seizure events.

#### Data collection

All experiments were performed in accordance to the guidelines and approval of Weill Cornell Medicine’s Institutional Animal Care and Use Committee.

#### Zebrafish Preparation

Mutant zebrafish larva (*Tg(HuC:h2b-GCaMP6f) x Tg(Vglut2a:dsRed)*) (24, 25, 52) were used at 5 - 8 days after fertilization - the h2b-tag localizes GCaMP to the nucleus to disambiguate the site of origin of the signal by eliminating contamination from axonal and dendritic sources. After anesthetizing for 1 min in 0.1 g/L tricaine-methanesulfonate (MS222, VWR TCT0941-025G), zebrafish were mounted dorsal up in a 35 mm petri dish (35–1008, BD Falcon) lid containing a bed of 1.5 ml hardened 1% agarose (Invitrogen 15510-027) by embedding with droplets of 1.7% low temperature agarose (Sigma A0701-100G). Once the agarose hardened, a small incision in the skin was made (typically) above the neuropil in the right optic tectum for electrophysiology (see below). The agarose around the tail of the zebrafish was removed to monitor tail movement. The larva was then paralyzed by point incubation in droplets of 300 μM pancuronium bromide (Sigma P1918-10MG) and covered in 2.5-3 ml of glucose-free 10% (vol/vol) Evans medium (2.9 mM KCl, 1.2 mM MgCl2, 2.1 mM CaCl2, 134 mM NaCl, and 10mM HEPES, pH 7.8) once tail movement ceased Epileptiform activity was induced with bath application of 15 mM pentylenetetrazol (PTZ) (Sigma P6500-50G), which is more common and effective than alternatives such as pilocarpine (22) and 4-Aminopyridine (26). Recordings were performed within 2 hours after embedding zebrafish in agarose.

#### Electrophysiology

Local-Field Potential (LFP) recordings were acquired with metal microelectrodes during initial seizure characterization. One ~2-3 MΩ tungsten microelectrode (WE30013.0F3, Microprobes) was inserted in the optic tectum record LFP and a silver wire was placed in the Evans medium to server as a reference. In some experiments, a second tungsten microelectrode was placed in either telencephalon or hindbrain region to confirm the global synchronous of ictal events. Electrophysiological recordings were amplified 1000x with a microelectrode amplifier (Model 1800, A-M Systems, Carlsborg, WA), digitized at 10K Hz using CED Power 1401, and stored with a computer runningSpike2 software (Cambridge Electronic Design).

During calcium imaging, the LFP was recorded using a glass microcapillary to avoid obstructing fluorescence excitation and light collection. Capillaries (1B120F-4, WPI) were pulled to a fine tip with vertical puller (P30, Sutter Instruments) to a tip diameter of 2-3 μm. The capillaries were filled with 4M NaCl after front-loading molten agarose (1.7% Ultrapure Agarose, Thermo) to the tip to prevent diffusion of the electrode solution once inserted into the brain. When tested in Evans medium, these electrodes had a resistance of 1-3 MΩ. The electrode was inserted through incision in the skin into the neuropil of the right optic tectum. Recordings were made with a microelectrode amplifier (Model 1800, A-M Systems, Carlsborg, WA) using bandpass filtering (between 0.1 Hz and 1 kHz) at 10 kHz sampling rate and digitized using custom MATLAB code.

#### Two-photon Imaging

A custom-built two photon microscope was used for imaging calcium activity and fluorescence in the embedded zebrafish. A MaiTai HP laser (Newport) was used to focus a 915 nm light beam through a water immersion 40x/0.8 NA objective (OBJ1; LUMPlanFl40XW/IR2, Olympus) onto the fish. Multiple separate horizontal planes of the fish were acquired sequentially in the dark at 2 Hz (256×256 pixels, 550×550 μm). The emitted light from the zebrafish larva was collected by the objective and directed towards two dichroic mirrors (DM1, 720dcxruv; DM2, 565dcxr; Chroma Technology) which reflected this light into two separate channels of green (F3; ET525\50m-2p; Chroma) and red (F2; ET605\70m-2p, Chroma) to be then amplified by the photomultiplier tubes (PMTs, R3896; Hamamatsu). A substage photodetector (PD, PDA36A, Thorlabs) detected light scattered from the focal plane and together with signal from the PMTs was processed by MATLAB based software ScanImage (53) to generate fluorescence images.

#### Analysis

##### Electrophysiology

Electrographic data was used to validate and examine ictal events during 2-photon imaging. The spectrogram of the LFP was calculated using short-time Fourier transform (*spectrogram* in MATLAB), with a window width of 0.1 s. Time of onset in the epileptiform events was determined using a statistical method based on the mean and 3 times the standard deviation (SD) of the power between 1-60 Hz during a 5 s interictal window. For electrographic analyses (**Fig.1**, **Fig. S1**), the first increase in power > 3 SD above the baseline was established as the onset of the event. The termination was set at the moment the LFP power returned within 3 SD of baseline.

#### Image analysis

##### Preprocessing

Images were collected by ScanImage and stored in tiff format. These data files were subsequently de-interleaved in Fiji (54) to extract the green (GCaMP) and red (DsRed) channels. Rigid motion correction was applied in MATLAB 2018a using the NoRMCorre algorithm and toolbox (55) on the red channel images, which did not contain activity-dependent fluorescence. The transformation matrix calculated on the red channel in each fish imaging series was then applied to the green GCaMP6f channel, yielding two motion-corrected tiff image series.

We used these data sets to examine ictal event activity across brain regions with two overarching analysis methods: First, we used pixelwise methods to characterize ictal event initiation, spread, and involvement across brain regions. Second, we performed single-cell analyses to observe ictal event development simultaneously in VGlut2a excitatory neurons and putative inhibitory interneurons, paying particular attention to E:I ratio and differential involvement of these two cell classes as well as network-level differences between them.

##### Correlation Coefficient Masks

During ictal events, GCaMP6f fluorescence in the image increases as the ictal event initiates and propagates through the brain. We first extracted relevant brain areas by using the preprocessed GCaMP6f data to calculate pixelwise Pearson’s correlation with the whole image average activity in the green GCaMP6f channel. To achieve this, we initially performed 2-dimensional Gaussian filtering on the imaging data (σ=2 pixels, ~4 micrometers) in MATLAB (*imgaussfilt*)and then smoothed individual pixels over time using a lowpass Butterworth filter (1^st^ order, cutoff frequency of .2 Hz). These data are hereafter referred to as “smoothed images”.

The individual pixel traces were then compared with the average of all pixel values in the image and a cutoff of .6 correlation (Pearson’s *r*) was used to remove any pixels not sufficiently correlated with the average image activity. To remove any remaining noise, any smoothed pixels not connected in groups of at least 80 smoothed pixels were removed. The resulting image mask, called the correlation coefficient mask, simply represents the extent of brain that was involved in seizure events over individual files. This mask was applied in subsequent analyses of ictal event development.

##### Lag analyses for individual ictal events

Individual ictal events were next detected in smoothed image series with a semi-automated method. First, general ictal event identification was performed: GCaMP6f fluorescence change was calculated as the frame-by-frame difference in average fluorescence, and peaks were extracted with a minimum peak height of average plus 2 SD. A minimum peak-to-peak distance of 8 seconds was also included. Due to slight variations in noise or atypically spreading ictal events in some data sets the standard deviation cutoff was adjusted or performed on the average fluorescence activity instead of the difference. When average activity was used to find peaks, we then used 10 sec windows around those extracted times to find the peak first derivative activity. We note that these adjustments to the automated process are only included to produce accurate windows for finding ictal onsets and would not change the onset times. This general identification produced the time at which the ictal event was increasing most rapidly in the fluorescence data. All individual ictal events were verified by visual examination of both the average GCaMP6f activity and the GCaMP6f image series videos.

Next, we used these ictal event times to create a window of whole-image ictal event activity of 6 seconds prior to this point, called the global template. This ictal event onset template was then compared with all individual pixels from the correlation coefficient mask in time windows around the individual ictal events: a 16 s window was used, starting at 10 s prior to peak and lasting 6 s past peak, a large time window that always captured the ictal event onset.

Then, the time of maximum correlation between the pixel trace and global template was found for each pixel from the mask yielding a matrix, for each ictal event, of the time when each pixel was most correlated to the global ictal event template. Pixels not significantly correlated (Pearson’s *r*, p<.05) with the ictal event average or with a peak correlation below *r*=.6 were excluded. When visualized as a heatmap this matrix provides a “lag map” of where the ictal event activity began and spread across the imaging field of view. To remove any remaining noise, these lag maps were subsequently smoothed by a 2-D Gaussian kernel (σ=6 pixels, ~11 micrometers). To account for the line-scanning nature of our 2-photon images, a temporal transformation matrix was applied to shift every pixel by its true time in milliseconds (e.g., the first pixel of a single tiff image was scanned 256 ms prior to the pixel in the middle of the image). Finally, when displayed, heat color indicates time relative to the average onset time calculated across pixels, negative values indicating pixels that were active earlier, positive values indicating pixels that were active later. A total of 10 separate animals produced N=177 ictal events.

From these lag maps, we used MATLAB’s edge detection functions to extract the regions of the fish brain that were ictal event-active at the earliest 500 ms. We also applied an area requirement of 300 connected pixels (~.5% of the 256×256 images) to exclude noise. These regions were used to measure which anatomical brain areas were most commonly invaded first by the ictal events. These regions were labelled the “initiation sites” in later single cell analyses. Other regions of the brain that were invaded later by the seizure are referred to as “propagation sites”.

To examine brain region involvement over different data sets, we compared the edges of initiation sites, per event, with the anatomical fish images (average fluorescence) and recorded the amount of times that an initiation site contained a particular brain region, given that the brain region was present (e.g., if the habenula was not visible in a specific ictal series then it was not included in the analyses of early onsets).

Correlations of these maps within fish were calculated as Pearson’s *r* (across seizures within the same FOV) to measure similarity of seizure propagation within individual animals. These correlations were thus only calculated when more than one event occurred in a particular FOV within a fish.

A summary map (**Fig. 3D**) was created within an individual fish by setting a temporal cutoff (500 ms) for early activity in the lag map, creating a binary mask from this cutoff, and then averaging these binary masks over different ictal events. Thus, if a pixel is involved in 100% of the ictal events of one fish then it will receive a 1, if a pixel is involved in half of the ictal events it will receive a .5, etc. The correlation matrix (**Fig. 3E**) was calculated on ictal series with multiple seizures in FOVs containing all listed brain regions.

##### Single cell analysis

Single cells were extracted from the unsmoothed GCaMP6f images using an automated custom MATLAB algorithm with an added optional step that allowed for manually marking any single cells missed by our algorithm. First, within a single data set, the lowest quartile of GCaMP6f activity frames were used to create an average GCaMP6f image. Background subtraction was performed using structural element disks in MATLAB (with *imadjust*) and then an algorithm was used to iteratively step through all pixel luminance values (0 to 255) down to the lowest 20% and extract all connected regions containing at least 4 pixels (~2 microns/pixel), but removing any cells containing more than 40 pixels. All images were visually examined and any missed cells that could be visually determined were added manually in MATLAB with a custom script (using *imfreehand*), but typically fewer than 5 cells were added at this step. Finally, we applied a correlation threshold cutoff to remove any single cells whose activity was not at least *r*=.3 correlated with the average activity of all the extracted cells. On average, our analyses extracted 351 seizure-active cells per image following these steps.

In separate cellular composition analyses, where we were interested in examining any detectable cells without requiring that they be seizure-active, we followed the above methods but used a lower Pearson’s cutoff of *r*=.1, which was low enough to include more cells than the *r*=.3 cutoff but high enough to exclude artifacts (such as occur on skin edges and near the animal’s eyes).

To separate putative inhibitory (expressing only GCaMP) and excitatory cell types (co-expressing DsRed and GCaMP in VGlut2.1+ cells) we took the red intensity values from individual cells and plotted their histogram. These histograms are characterized by two peaks: the first peak, lower red values, denotes the GCaMP-positive cells that do not co-express DsRed (putative inhibitory interneurons); the second peak, higher red intensity values, denotes the GCaMP-positive cells that also co-express DsRed (putative excitatory neurons). These histograms were fit by a dual gaussian mixture model, and the mean and standard deviation of both component gaussians were extracted (*fitgmdist* in MATLAB). The cutoff used in our study for labeling a cell as excitatory was two standard deviations past the first gaussian’s mean. In some cases, artifacts from the zebrafish skin, eyes, or a tilt in the animal’s body could affect these histograms. In those cases, the histogram fitting was performed regionally by manually selecting anatomical regions (applying MATLAB’s *imfreehand*) and then performing the dual gaussian fit. All images were visually examined after cell type dissociation by plotting cell types on composite red/green images of the larval zebrafish to verify appropriate labelling by the algorithm. In two data sets we could not extract E vs I cell type differences (due to either imaging or fluorophore labelling technical difficulties), so the 10 ictal events in those data series were not included in our E:I single cell analyses leaving N=167 ictal events.

###### Seizure onsets

In order to compare results across animals and seizure events, we extracted ictal onset times from each ictal event for all single cells. General ictal events were found above (see Lag Analyses), but for single cell analyses we sought to determine the exact time of cell onsets relative to a population onset. Thus, we applied a similar correlation method as the pixelwise lag analyses above but with global templates that were centered around the peak second derivative of single cell population activity, the time of fastest recruitment of individual neurons to a seizure event.

Single cell involvement in seizures was determined by identifying the times at which individual cells became significantly involved in the ictal events (**Fig. S4**). To find these times in single neurons we filtered the cell GCaMP6f profiles by temporally smoothing in 2 s windows (boxcar average) and then compared these traces in time windows around ictal event onsets that were found in the lag analyses above.

Global response templates in this analysis were created by taking the average cell response from a period of equal length between the first and second peak derivative of the average single cell fluorescence traces. This created a template centered on the peak acceleration point of the ictal event, a time that more accurately represents true ictal event onset compared to other methods of cutoffs such as half-height or 2-standard deviations above the mean. We refer to this peak acceleration time of the population activity as the ictal onset.

Next, we found the peak linear correlation (Pearson’s *r*) between this template and individual cell fluorescence traces in a window from 8 s before and 3 s after the period of the global template. This wide temporal window was important because our method of global template creation depends on the rise time of the ictal event: slowly evolving ictal events will have longer templates than faster events. (Note: the correlation window used for single cell onsets was 11 seconds plus the length of the global template, making it a window approximately equal to the 16 sec window used in lag analyses above). Our algorithm required that a single cell be at least *r*=.8 correlated with the global template to be included as having an onset in the ictal event, a strict requirement to reduce the chance of erroneous single cell onset classifications. The onset time for a single cell was classified as the time where it reached 5% of peak correlation (above the .8 cutoff) with the global template. This 5% requirement was included because many cells displayed wide peaks of correlation with the global template: taking the earliest peak correlation most accurately represented the onset of a single cell in the ictal event. Because some cells displayed small fluctuations in fluorescence that coincided with ictal onsets, when cells exhibited more than one *r*=.8 peak during the time window analyzed, we took the later *r*=.8 peak to prevent erroneous early-onset classifications. Finally, all individual cell onsets were temporally adjusted for the 2-photon system’s scanning (as done for the pixelwise lag analysis, shifting by the pixel scan time from the cell centroid), and the population activity was temporally corrected by 256 ms (the midpoint time of the frame scan). This method was applied to all data sets and all cells regardless of E or I classification.

When temporally aligning data we used the average ictal onset time calculated across all cells significantly recruited to each individual seizure, a procedure analogous to the relative times calculated in our lag analyses above. Thus, data are displayed relative to this time zero point.

Inter-ictal periods were defined as all times outside of ictal events (after PTZ application). We classified the termination of ictal events as the point at which the average fluorescence activity returned to its value at the second derivative onset, which was used to separate ictal vs interictal time periods for later analyses.

###### Excitation:Inhibition (E:I) balance

We examined E:I relationships by calculating an E:I ratio index. We use this method instead of simple E/I ratio because in some cases we observed low numbers of cells in our initiation sites. E:I index values in initiation regions were calculated by summing these cell types within the earliest active brain areas (see above, Lag analyses). E:I ratios in propagation zones were calculated the same way, using remaining areas of the brain that were significantly involved in seizures (propagation zones). The index value was calculated as

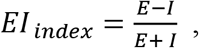

where E and I are counts of E and I cells. The E:I ratio index provides a measure of the relative differences in amounts of E and I cells located in the initiation and propagation regions.

We measured excitatory and inhibitory cell recruitment throughout ictal events by temporally aligning all events and then calculating the fraction of E and I cells significantly involved in the ictal events before and after the onset.

##### Ictal vs interictal brain state and network analyses

###### Dimensionality metric

A population of N neurons makes an activity trajectory in the N dimensional space over some time points (T). The effective dimensionality of the trajectory can be quantified with the uniformity of the singular values (λ_j_) of the trajectory matrix (NxT, mean-subtracted).

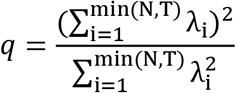

If the singular values are perfectly uniform, all principal components explain equal amount of variance, and therefore the effective dimensionality should maximal. If only one singular value is non-zero, then the effective dimensionality should be 1. For our dimensionality analysis, the number of cells (N) was variable but always greater than the number of time points (T=10s*2Hz=20 samples) of our time window, so the upper limit of the dimensionality was fixed at T. This allowed us to compare the effective dimensionality across trials with different number of cells without further adjustments. A dimensionality at an instantaneous time-point t is estimated by calculating the dimensionality over the time window centered around t. The instantaneous estimation was used to construct a time-series from which we could visualize the time evolution of dimensionality.

###### Excitatory & inhibitory correlations

Populations of excitatory cell pairs (e-e) and inhibitory cell pairs (i-i) were analyzed for their correlations around ictal events. The correlation metric (*C*) is an average pairwise correlation value of the calcium traces over a time window (1.5 s),

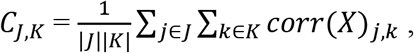

where X is a matrix whose rows are a time series of a single cell within the rolling time window, and J and K are sets of indices for excitatory or inhibitory cells. *corr*(X) returns a correlation matrix. Self-correlations (diagonal elements) are ignored. A correlation at an instantaneous time-point t is estimated by calculating the correlation over the time window centered around t. The instantaneous estimation was used to construct a time-series from which we could visualize the time evolution of correlation.

##### Statistical Analysis

We applied different statistical tests on our data as appropriate: paired comparisons were performed with the Wilcoxon signed-rank test, and unpaired comparisons were measured with the Wilcoxon rank-sum test, both of which are non-parametric tests. The significance levels and N values are described whenever the tests are applied in the Results.

## Acknowledgments

The authors thank Catherine Schevon, Michael Wenzel, Jyun-you Liou, and Jonathon Victor as well as members of the Aksay and Schwartz labs for constructive discussion of this manuscript.

## Author contributions

Conceptualization: H.M., E.A., and T.H.S. Methodology: P.G., S.S., and H.M. performed data acquisition and J.E.N., C.C., and H.M. performed data analysis. Investigation: P.G., S.S., and H.M. performed data acquisition and J.E.N., C.C., and H.M. performed analysis. Visualization: J.E.N., C.C., J.L. Supervision: H.M., E.A., T.H.S. Writing: J.E.N. and E.A. wrote the original draft and J.E.N., E.A., and T.H.S performed review and editing.

## Data availability

Data and code needed to evaluate conclusions of this paper will be uploaded to the Zenodo data repository.

## Supplementary Materials

**Figure S1.**
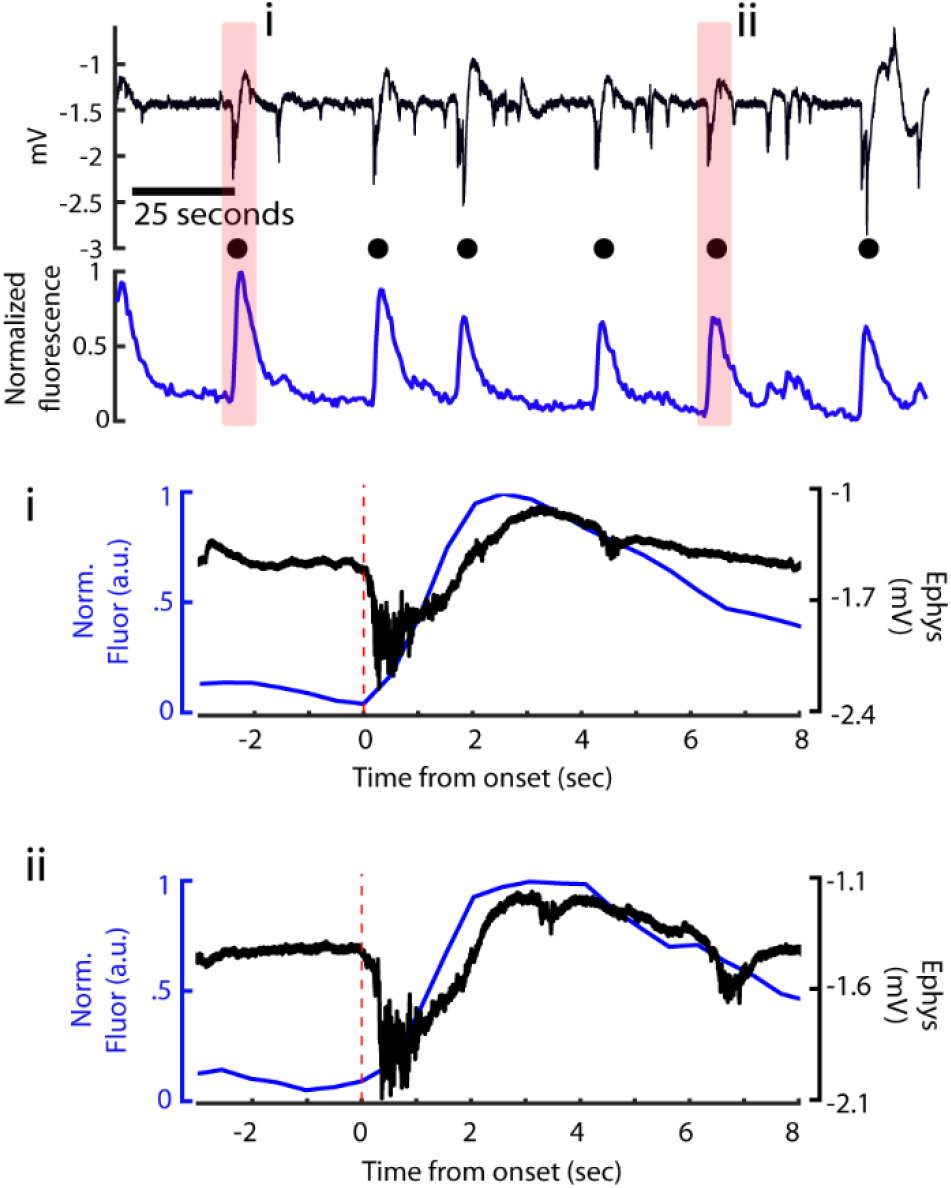
Identical timing of ictal events in calcium and electrophysiological signals. A sample series of ictal events is shown from **Figure 1**, with two seizures highlighted (pink). The first event is shown in higher temporal magnification (i), as well as the fifth event (ii). A 2 Hz notch filter was applied to the electrophysiological signals in the bottom panels, and the calcium activity is taken from the entire imaging frame. Our onset detection method was applied to all seizures (red dashed line).

**Figure S2.**
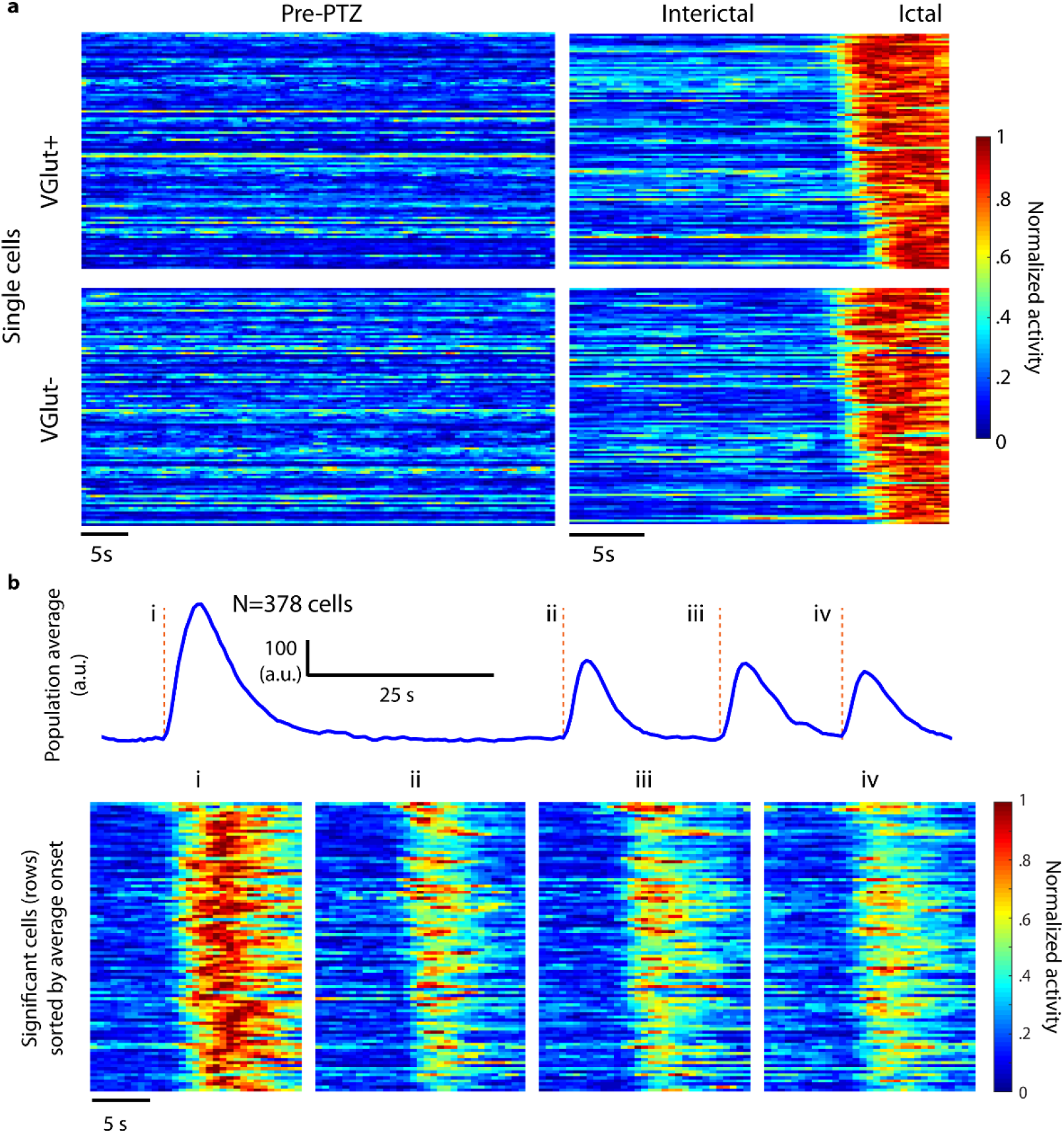
Single cell activity. (**A**) Individual cell activity is shown (rows) for the same cells prior to PTZ administration (*left*) and interictal activity prior to an ictal event (*right*) after PTZ administration. Excitatory and inhibitory cell types are shown separately and sorted by detected onset time within this ictal event. (**B**) Top: cell population activity demonstrates a series of four ictal events; Bottom: single cell activity during these events is shown sorted by average onset across events.

**Figure S3.**
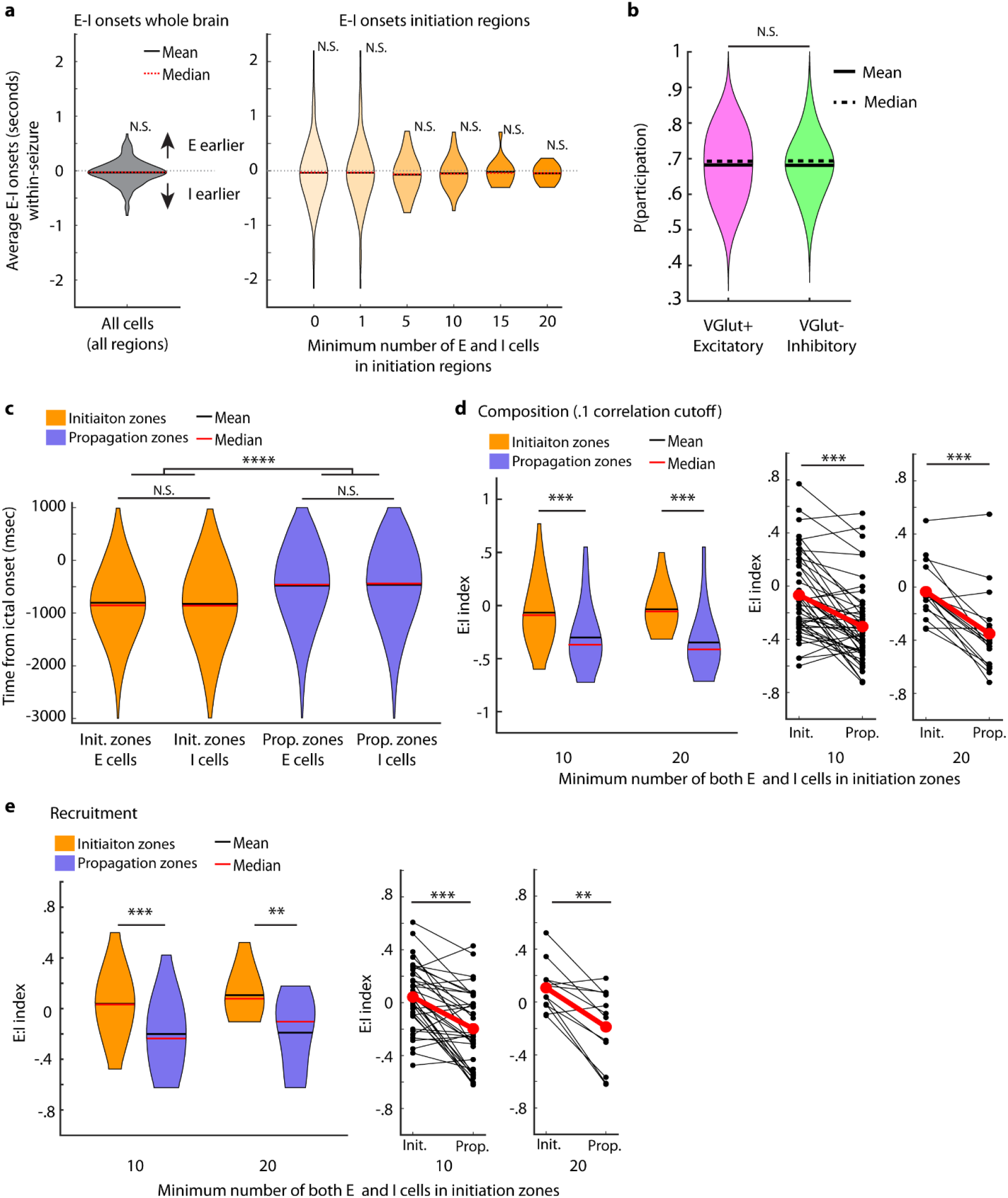
Associated with Figure 4 E:I and onset timing. (**A**) Left: the difference in average onset time between excitatory and inhibitory cell populations is shown (positive values indicate earlier excitatory cell onset) for whole brain Right: same analysis for initiation zones. Initiation zones are divided into varying sizes (All seven violins: N=167, N=167, N=101, N=62, N=36, N=18, N=12 ictal events; none were significant by α=.05, Wilcoxon signed-rank test). (**B**) Probability of participation was calculated across all E and I cells that participated in more than one seizure (N=40 data series with at least 2 seizures, 159 seizures total; N.S. α=.05, Wilcoxon-rank-sum test). (**C**) Recruitment times between E and I cell types (N=1264, N=1135, N=12143, N=14992; N.S., α=.05, Wilcoxon rank-sum test) and between initiation and propagation sites (**** p<.0001, Wilcoxon signed-rank test). (**D**) Number of E and I cells detected in regions that were determined to be initiation and propagation zones, regardless of whether these cells participated in seizure events, shown across multiple size cutoffs (N=49, N=18, Wilcoxon signed-rank test, *** p<.001). For all data (167 seizures), with no cell count cutoffs, E:I composition in initiation regions was −0.069 +/- 0.042 (SEM), while E:I composition in propagation regions −0.232 +/- 0.024, corresponding to a roughly 29% increase in E:I ratio in the initiation zone relative to the propagation zone across all seizures. (**E**) Number of E and I cells recruited in initiation and propagation zones shown across multiple size cutoffs (N=36, N=12, p<.001, Wilcoxon signed-rank test, ** p<.01, *** p<.001). For all data (167 seizures), with no cell count cutoffs, E:I recruitment in initiation regions was 0.034 +/- 0.043 (SEM), while E:I recruitment in propagation regions was −0.162 +/- 0.022, a significant difference corresponding to a roughly 40% increase in E:I ratio in the initiation zone relative to the propagation zone across all seizures.

**Figure S4.**
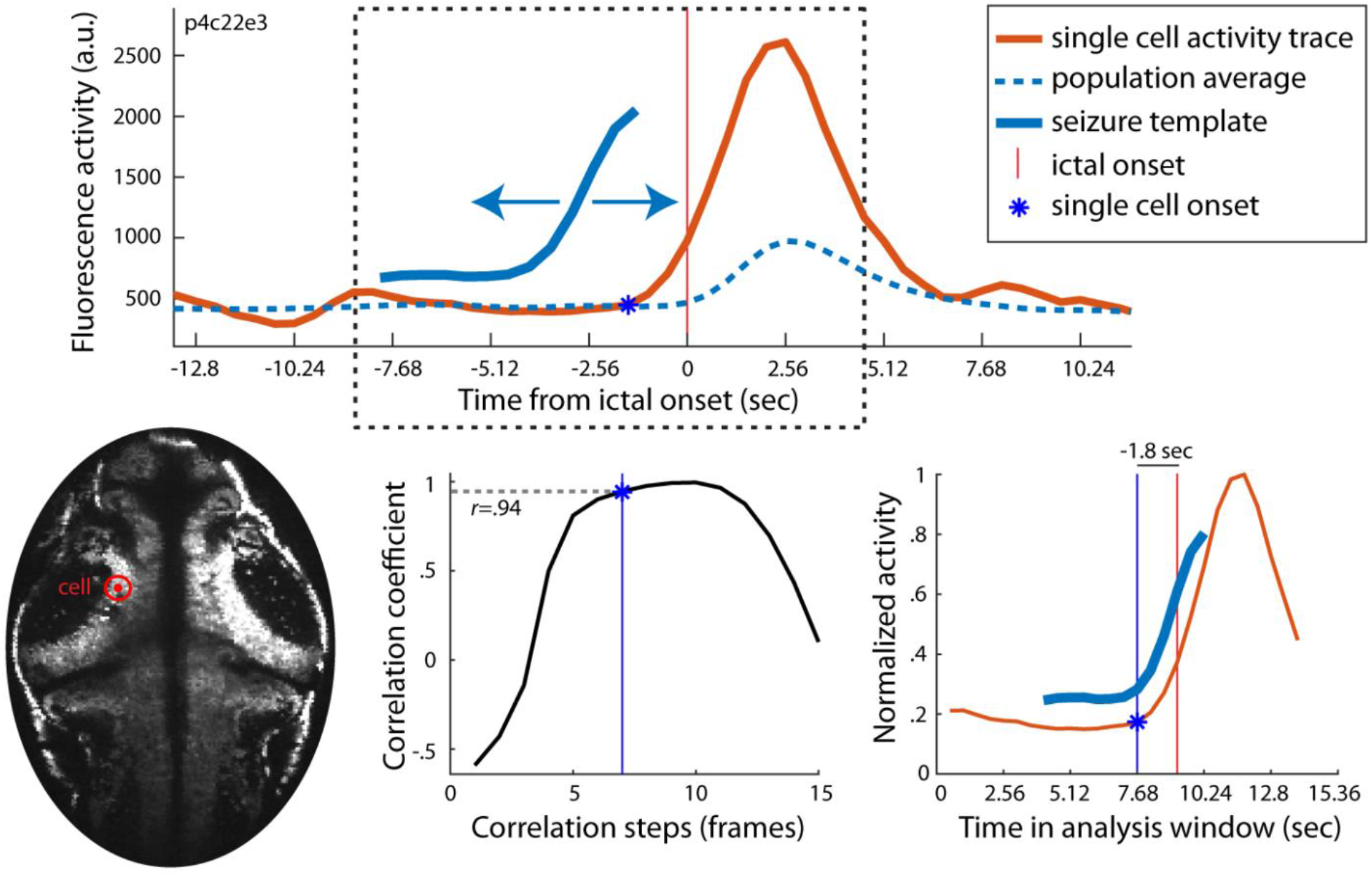
Single cell onset detection. Ictal events were detected from global (all cells) activity, and a template of global activity was created around the peak second derivative of this trace. Top: this template was slid along single cell activity traces to determine the estimated onset of the single cell based on its correlation with the global template (see Methods). Bottom, left to right: the single cell location in the fish, the correlation at each step of the analysis with the best fit location marked, and the template aligned with the single cell onset estimate. This single cell’s onset was ~1.8 seconds earlier than the population ictal onset.

